# Cortical spike multiplexing using gamma frequency latencies

**DOI:** 10.1101/313320

**Authors:** Dana H. Ballard, Ruohan Zhang

## Abstract

One of the fundamental problems in understanding the brain, in particular the cerebral cortex, is that we only have a partial understanding of the basic communication protocols that underlie signal transmission. This makes it difficult to interpret the significance of particular phenomena such as basic firing patterns and oscillations at different frequencies. There are, of course, useful models. Motivated by single-cell recording technology, Poisson statistics of cortical action potentials have long been a basic component in models of signal representation in the cortex. However, it is increasingly difficult to integrate Poisson spiking with spike timing signals in the gamma frequency spectrum. A potential way forward is being sparked by new technologies that allow the exploration of very low-level communication strategies. Specifically, the voltage potential of a cell’s soma now can be recorded with very high fidelity in vivo, allowing correlation of its fine structure to be correlated with behaviors. To interpret this data, we have developed a unified model (gamma spike multiplexing, or GSM) wherein a cell’s somatic gamma frequencies can modulate the generation of action potentials. Such spikes can be seen as the basis for a general-purpose method of modulating fast communication in cortical networks. In particular, the model has several important advantages over traditional formalisms: 1) It allows multiple, independent processes to run in parallel, greatly increasing the processing capability of the cortex 2) Its processing speed is 10^2^ to 10^3^ times faster than population coding methods 3) Its processes are not bound to specific locations, but migrate across cortical cells as a function of time, facilitating the maintenance of cortical cell calibration.

## 1 Introduction

The action potential was described by Bernstein [1902], but it was not until 1952 that its underlying conductance-based mechanism was modeled by Hodgkin and Huxley [Hodgkin and Huxley, 1952]. Having a basic model of action potential generation opened up the study of the meaning of sequences of such potentials (spikes). Spikes are used in many specialized ways in different parts of the brain, from the rate codes in the brainstem’s vestibulo-ocular reflex circuitry [Highstein et al., 2005] to the tonically-active signaling in the basal ganglia [Graybiel et al., 1994] and phase coding in the hippocampus [Skaggs and McNaughton, 1996, Sreenivasan and Fiete, 2011]. In contrast, the basic codes used to model spikes in the cortex, while of great usefulness as a correlate of behavior, have so far eluded an understanding that is completely satisfactory.

The most basic firing pattern of cortical cells exhibits Poisson distributions [Shadlen and Newsome, 1998, Gerstein and Mandelbrot, 1964, Berry II and Meister, 1998]. These distributions, which comprise the core of various population coding models, including rate coding [Gerstner et al., 1997], have allowed the interpretation of thousands of experiments. However any literal realization of spikes in population coding is expensive. When all the costs are taken into account, spike generation and transmission consumes on the order of 50% of a cell’s metabolism [Lennie, 2003]. Population coding also imposes demands on the necessary number of underlying neural synapses. A striate cortex simple cell model receiving feed forward input from a 10 × 10 image patch could require ~ 10^4^ synapses, a figure on the order of the total number of synapses estimated for a neuron [Schüz and Palm, 1989].

To add to these difficulties, there is the challenge of using Poisson spikes to implement multiple independent processes. This conundrum was originally characterized as the ‘binding problem,’ specifying the co-reference of an object’s cortically distributed features [Von der Malsburg, 1999, Singer, 1999a, Seymour et al., 2009], but a more recent focus from several perspectives is that of managing temporally coextensive *processes*, as shown by the following examples. Alternate motor plans may be active prior to choosing between them [Cisek, 2012]. Analyses of cortical recordings suggest that spikes may represent mixtures of multiple task features [Kobak et al., 2016]. The hurdle in addressing this multiplexing capability is in specifying a way to dynamically partition the brain’s extensively connected networks in a way that they do not interfere with each other.

One approach to achieving multiprocessing is to take advantage of spike timing. The classic spike timing treatise is by Abeles [1991], its fundamental observation being that the arrival of synchronous inputs facilitates action potential generation [Diesmann et al., 1999, Pfister and Gerstner, 2006]. Consequently, several timing-based interpretations of neural coding have been suggested [Diesmann et al., 1999, Pfister and Gerstner, 2006, Dettner et al., 2016, VanRullen and Thorpe, 2002]. However, recent experimental evidence suggests that the most likely timing approach used by the cortex is to exploit the gamma frequency band in the range of 30 to 80 Hz. Increasing numbers of experiments exploring network behavior at this frequency band [Buzsáki and Wang, 2012, Gray et al., 1989, Engel et al., 1999, Singer, 1999b] are starting to be functionally interpreted [Womelsdorf et al., 2006, Fries et al., 2007, Landau and Fries, 2012, Bastos et al., 2012]. The major function of the gamma band is a source of modulatory signals used in large-scale network communication [Fries et al., 2001, Chalk et al., 2010, Fries, 2015]. The gamma band encompasses many associated computational issues, but for purposes of discussion, they can be divided into those related to the use of the band as a *medium of communication*, in Fries’s words ‘communication through coherence,’ and those relating to *messages sent* using features of the band. The former focus can also include larger process control issues that involve theta, alpha and beta frequencies [Von Stein and Sarnthein, 2000, Fries, 2015], which we do not cover herein.

The focus of the paper is neural network multiplexing. Without the ability of running multiple simultaneous processes, the brain must solve the problem of initiating and terminating single processes in sequence, with the attendant temporal overhead, a very unlikely prospect. This implausibility leaves the option of parallelism, one form of which uses multiplexing where individual neurons can somehow manage spikes from different processes. That the cortex might use some kind of mixing is gaining credence [Pillow et al., 2005, Kobak et al., 2016]. The paper describes a model termed ‘gamma spike multiplexing,’ or GSM, that defines such spike mixing networks, and further shows that they can be set up in a way that allows circuits’ effective connectivity to randomly migrate in the course of carrying out their computations. The central representational feature of the model takes advantage of the over-completeness^1^ of cortical maps to select subsets of neurons that represent the state of a computation probabilistically on each gamma frequency cycle. Consequently, the computation for any particular problem the cortex is attempting to solve can be represented by different neurons on each gamma cycle. Simulations demonstrate that the multiplexing process allows on the order of 4 to 16 simultaneous separate computations to be mixed successfully among the same neurons with little or no cross-talk. This mechanism opens up the possibility that multiple independent low-level routines may be carried out in parallel. Furthermore the mixing process can still result in individual neurons exhibiting classical Poisson statistics, making it compatible with conventional single cell recording data.

## 2 The model

The GSM model borrows elements from other research, but features two innovative central interlocking components, each of which are advances over Ballard and Jehee [2011]. The first is a way of isolating parts of a network so that it can implement a specific neural computation. This capability is satisfied by assuming that, for the duration of the computation, time-sensitive subsets of neurons are responsive to a specific gamma frequency. Multiplexing behavior can be implemented very economically if the gamma frequency on a cell’s soma can be randomly modulated between a small discrete set of specific gamma values.

The second component is that from a pool of neurons that could represent a computation, a random subset are selected on each gamma cycle. Thus, the subset of neurons representing any computation is different from cycle to cycle, such that a computation is not represented at any specific location, but migrates throughout the network. In the case of more than one active computation, each one needs to have its own gamma frequency. Combining the different sets of gamma frequencies with the random selection of cells, means that all of the computations can migrate throughout the network and share neurons. The sparseness of action potentials results in very low probabilities of spike interference.

The following three subsections expand this summary as follows: 2.1 describes the management of multiple gamma frequencies in each cell’s soma, 2.2 describes the specification of the proposed mechanism for probabilistic cell selection, and finally 2.3 exhibits simulations of the model that show the interaction of multiple gamma frequencies and probabilistic cell selection.

### 2.1 Multiplexing via gamma modulation

At any one instant, a private subnetwork can be delineated by associating an appropriate set of neurons with a specific gamma frequency. The network has the property that a distinct subset of neurons is chosen probabilistically for each gamma cycle. Thus in space and time, the participating neurons are usually different and separated. However even though a single computation being carried out uses a different network on each cycle, every generated spike therein is clocked to a multiple of the chosen gamma frequency. This leaves open the question of how to fit the computational process onto this moving substrate, which will be tackled in subsection 2.2 for a specific example.

In the case of a single computation, its gamma frequency would be easily detectable by standard methods. However, this situation changes with the introduction of additional computations that use multiplexing. Such computations can share cells probabilistically, each with a separate frequency from the gamma range. The randomness in the selection of neurons results in *any* neuron being able to be used for different computations, each with spikes indexed with a unique temporally-restricted local frequency. This point requires some elaboration. If computations are going to share a cell, and each of these computations is associated with a different gamma frequency on its soma, the net result is the soma will be expected to modulate its gamma frequency in time.

This concept is illustrated artificially in Fig. 1 by taking a set of spikes from one cell in a simulation, and interpolating the time axis with gamma range frequencies. In an 800 millisecond interval, Fig. 1a shows a set of spikes that are associated with separate gamma modulating frequencies. Fig. 1b shows an artificially constructed modulation, stitching together particular modulating frequencies. While there is still a complete understanding of how the neural circuitry could control this functionality, frequency modulation has been observed via patch clamping [Atallah and Scanziani, 2009, Perrenoud et al., 2016]. The outline of one abstract possibility is shown via Fig.1c. The input to the cell’s dendrites consists of clusters of spikes realizing a delay code on the order of 5 *~* 10 milliseconds, and this is multiplicatively modulated by inputs proximal to the soma. In the GSM simulation of this process, the action potential generation component is simplified to the conventional normalized dot product **x · w**, where **x** represents the vector of input spike codes and **w** represents their recipient synapse strengths. The normalized dot product is converted to a delay as shown in Fig. 2a, which the simulation uses in its spike distribution calculations.

**Figure 1:**
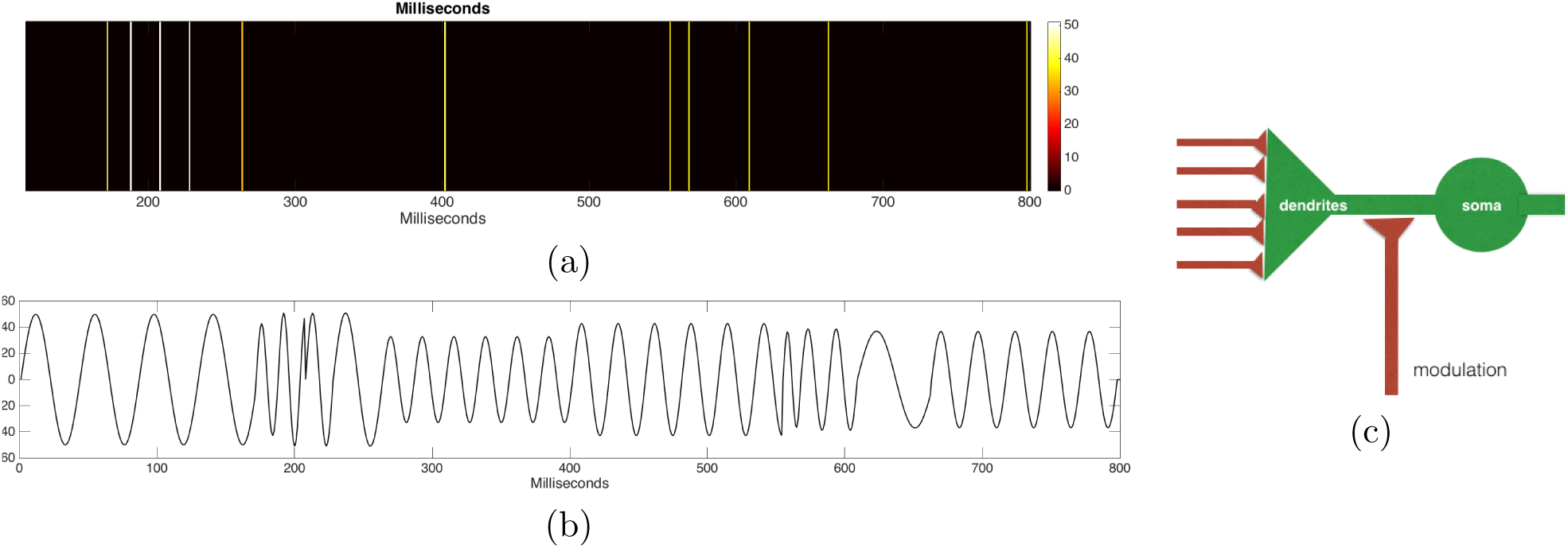
Parallel ‘wireless’ connection via simultaneous gamma modulation. An illustration from the simulation data from a single basis function that shows Gamma frequency modulation based on spike timing. (a) Generated spikes for different computations that are labeled by their associated gamma frequencies (See colormap in Hz). (b) Illustrative gamma modulation constructed to match their zero phase points to coincide with their corresponding spike. For any spike generation instant as long as all the neurons in the cohort are identically modulated, they will be effectively connected. (c) Schematic of the overall architecture posited for making it sensitive to spiking at a the rise point in the gamma cycle. Thus it acts as a filter for incoming spikes in that interval. Other neurons that are also identically modulated in that interval form apart of components of a desired interconnected network. If the somas of a large network are randomly modulate by a discrete set of frequencies, then subsets of neurons using instantaneous common frequencies define effectively private networks.

**Figure 2:**
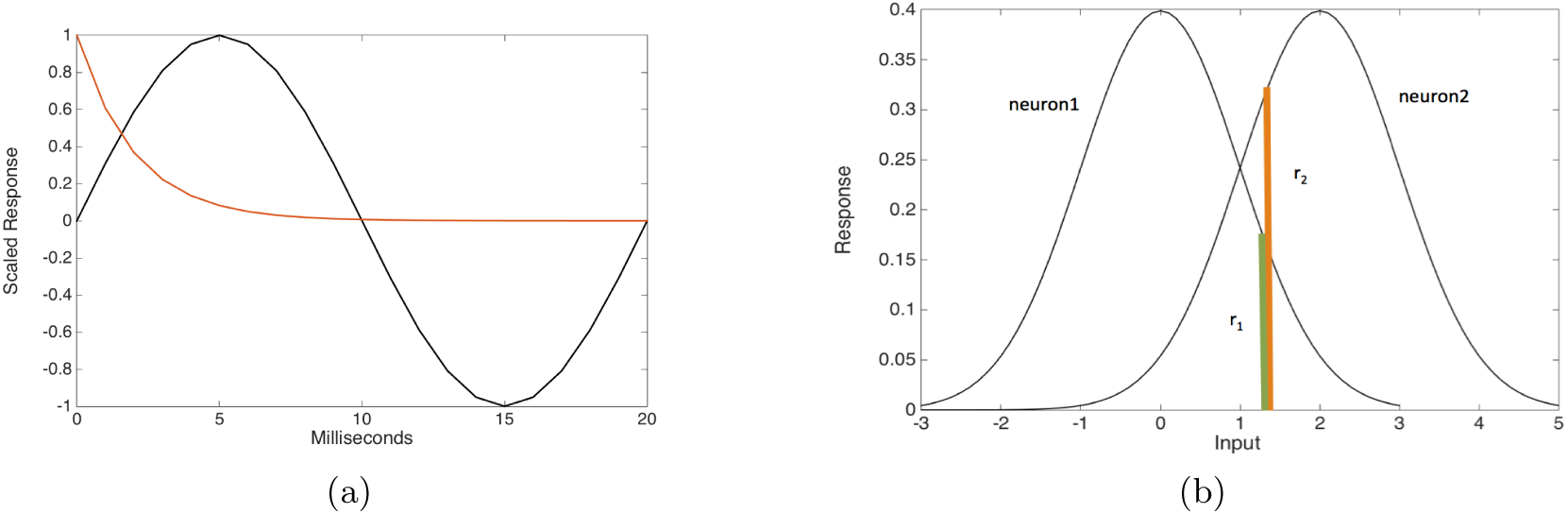
Methodology for computing spike codes within a single gamma period. (a) A spike signals an analog number coded as a phase delay from a gamma frequency zero crossing. The red curve shows response magnitude as a function of the delay. Short delays represent large magnitudes and long delays small magnitudes. The total range of delays is assumed to be on the order of 5 ~ 10 milliseconds. (b)To produce Bayesian statistics the basis functions have to be selected probabilistically. The relative magnitudes of the input’s projections determine the probabilities of a neuron being selected to be part of the representation of the input, e.g. the probability of basis function *r*_1_ being selected is 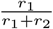.

Extending this temporal coordination to a network needs elaborate support circuitry to implement it successfully [Brunel and Wang, 2003, Atallah and Scanziani, 2009], but if these mechanisms are in place, a very elegant way of instantaneously connecting large networks is possible. Fig. 1 suggests a possible mechanism. Consider a time window long enough to represent several gamma cycles at a particular frequency. If these oscillations were co-modulated among a random subset *simultaneously* with all the neurons in that cohort would be engaged in a particular computation via using the same gamma frequency. Without any elaborate interconnecting circuitry, those neu-rons would behave as being effectively connected by virtue of being filters for spikes arriving in a narrow time window. All this structure might seem a little exotic, but data from patch clamp recordings [Perrenoud et al., 2016] reveal that the soma of cortical cells is continuously oscillating in the gamma range. Atallah and Scanziani [2009] have demonstrated that the needed gamma modulation can be controlled by adjusting excitation and inhibition. This demonstration is complemented by the circuit model of Brunel and Wang [2003], that illustrates these features and the additional possibility of *~* 10 ms spike phase lags.

### 2.2 Parallel probabilistic computation

In the model, each of the computations would be solving separate arbitrary problems. The implicit general assumption is any arbitrary process will be able to be expressed as an attractor dynamics with a time constant of a small amount of gamma cycles. However for illustrative purposes the situation is simplified in this exposition. Instead of arbitrary computations, we use multiple instances of the familiar computation of representing image patches with receptive fields in striate cortex using sparse coding. This setting is useful as it is familiar, yet lays bare several issues that would have to be tackled in any process encoding.

The sparse coding example, due to Olshausen and Field [1997], asserts that the early visual cortex prefers to develop neurons with receptive fields structured to represent the statistics of natural images, and moreover the receptive fields for such representations are chosen to have relatively few neurons active at any one time. This coding strategy can be achieved if the set of learned receptive fields is sparse, meaning that the number of chosen receptive fields is significantly less than the number needed to represent the input’s degrees of freedom exactly. Such a reduced set can be chosen for any particular input.

The GSM model assumes that a spike can represent an analog quantity as a phase delay [Vinck et al., 2010], with a reference point chosen as the gamma frequency’s zero crossing, as shown in Fig. 2a.

As Pouget et al. [2013a] point out in their review, “There is strong behavioral and physiological evidence that the brain both represents probability distributions and performs probabilistic inference.” When these distributions are distributed in networks, fast inference algorithms are possible that can complete the settings of the network from given sets of marginal probabilities Pearl [1986, 2014].

There has been a great amount of research into the neural representation of such probabilities in neural networks, but the large majority of them are subject to the earlier criticisms advanced of population coding. The GSM formalism posits that spikes signal explicit analog values at gamma frequency rates and consequentially computationally is much faster.

Regardless of the representation, a crucial feature in coding stimuli is that the neurons are *selected* probabilistically with odds chosen proportionally to their receptive field projections. This feature is dictated from that the constraints that arise from having to learn the receptive fields [Jehee et al., 2006b]. Fig. 2b shows an example for just two neurons. If the two responses are at levels *r*_1_ and *r*_2_ respectively, then their odds of being picked are in the ratio *r*_1_: *r*_2_. This method can then be generalized to however many neurons are in a larger pool.

Since the calibration of receptive fields is an ongoing process, selecting neurons in this way means that the overall distribution of the receptive fields obeys Bayesian statistics, e.g., [Jehee and Ballard, 2009, Jehee et al., 2006a] by ensuring that all receptive fields participate in the receptive field adjustment process appropriately. The reason for this is in learning the receptive fields in the first place, the cells must be sampled appropriately often for the proper receptive fields to form. By contrast, if maximum likelihood selection is used, a restricted portion of cells will form receptive fields at the expense of others, with the consequence that sampling such distorted fields for stimulus coding will not be Bayesian.

For sparse coding to be a candidate for the model, its algorithm must be able to compute its receptive fields in one parallel gamma step, whereas the standard method of computing receptive fields computes them sequentially. Parallelism can be realized if the number of receptive fields in the code is increased by a factor of 5 to 10 over conventional approaches. Fig. 3 compares the two approaches. Both the the input image (black) and cells’ synapses (gray) have dimensions *n*^2^, but can be depicted succinctly as vectors. Part(a) shows the sequential method. The contribution of a cell (red) selected for representing the image can be determined by subtraction (green). The residual is now a candidate for further representation by other cells. In contrast the parallel method in part (b) chooses 50 cells in parallel. Since they are each chosen by having positive projections onto the input vector, adding them up and normalizing the sum, results in a very good code for the input.

**Figure 3:**
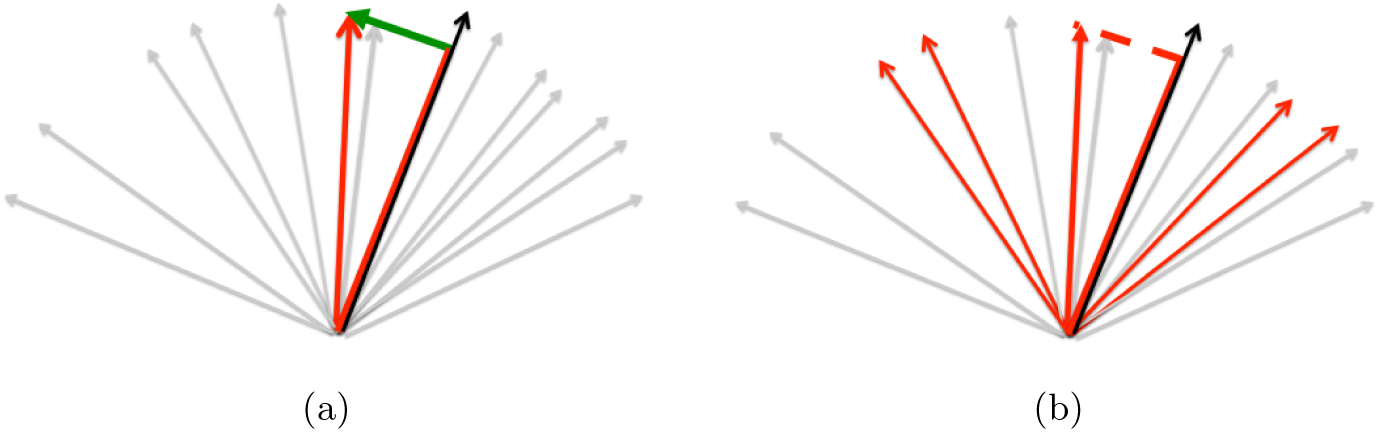
sequential and parallel sparse coding methods compared. (a) Standard algorithms for representing an input (black arrow) pick an approximating neuron (red) and sequentially fit the residual (green), a process that requires multiple iterations. (b) The GSM chooses 50 representing neurons in a single parallel step and then sums and normalizes the result.

Fig. 4a shows successive reconstructions using 5, 10, 20, 50 and 100 basis functions. If sets of cells of size 50 to 100 are used to code the input of of modest size image patches in parallel, the resultant code results in accurate approximations. Degradation is very apparent for the low number of basis functions reconstructions but this is to be expected given that the 14 × 14 image patches have 196 degrees of freedom. The accuracy of the representation improves as a function of the number of cells, as shown in Fig. 4b. However, note that even after the number of cells is sufficient to produce a useful representation, adding cells continues to improve the accuracy.

**Figure 4:**
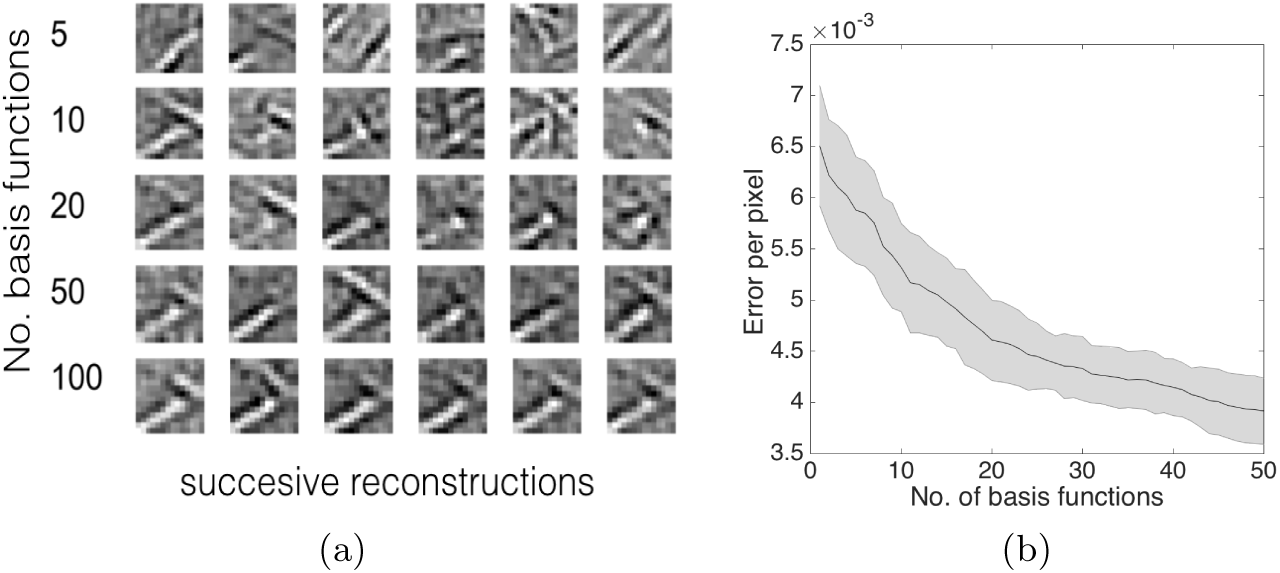
The parallel representation of the input improves its fidelity with increasing numbers of basis functions. (a) Sets of six independent reconstructions of a particular image patch using increasing numbers of basis functions, as indicated in the extreme left column. (b) Reconstruction error as a function of the number of basis functions used displayed with standard errors of the mean.

Although the example is specialized, its properties generalize in a very important way. The primary facets of this representation are 1) that the receptive fields can be computed in one parallel step; they can be selected randomly and 2) the random selection provides an essentially limitless number of combinations of cells. In the simulation, 50 were chosen at random from a library of 1000 cells repeatedly. Thus, the representation that codes a specific temporally invariant input wanders around the network differently on each gamma cycle.

### 2.3 Process Mixing Simulation

The simulation is straightforward. The initial step is to specify the number of processes. For each process, a start time, gamma frequency and duration(in terms of gamma cycles) are chosen. The simulation clock is advanced each millisecond. In the simulation if a cell is in use at a given it locks out other inputs. The simulation takes the shortcut of generating each process’s spikes sequentially. The assumption is that competing inputs that collide with a process’s input will be rare owing to the sparseness of spikes in the network, and if they do the perturbation to the signal will be small. In the same way if less than the selected number of cells is allowed the error will be likewise very small (See Fig. 4).

Table 1 shows the parameters chosen for the particular simulation. The central parameters chosen for the processes are chosen to be representative but are unknown in a mammalian system. In primates, the number of simultaneously active processes may be linked to working memory, and in that case the number would be four. However much less is known about the use of unconscious, over-learned memory, which may have a significantly higher figure. The individual gamma frequencies are chosen from the low gamma range and last for an average of 400 milliseconds. A process’s start time is random.

**Table 1:** Parameter values used in the simulations. In the simulation, for each process, its governing parameters are selected. Next, its basis functions (bfs) are selected probabilistically. If there is free space, spike delays computed on the basis of their projection values. The time for the delay and following refractory interval is marked as in use. *𝒩* (*m, σ*) is the normal distribution.

| Name | Value | Units | Description | Function |
| --- | --- | --- | --- | --- |
| $T$ | 800 | milliseconds | total length of simulation | simulation |
| $n_b$ | 1000 | scalar | total number of bfs | simulation |
| $n_s$ | $14 \times 14$ | pixels | size of input & bfs | simulation |
| $n_r$ | 50 | scalar | number of bfs to represent the stimulus | simulation |
| $N_p$ | 16 | scalar | number of processes | process |
| $f_p$ | $35 + \text{rand}(1, 20)$ | Hz | frequency to modulate a process | process |
| $t_\gamma$ | $20 + \mathcal{N}(0, 2)$ | scalar | process duration in gamma cycles | process |
| $t_s$ | $\text{rand}(1, T - \Delta)$ | milliseconds | process start time(Not too close to the end) | process |
| $d_\gamma$ | $T_\gamma/2$ | milliseconds | max length of the delay | spike |
| $d_r$ | 3 | milliseconds | refractory period | spike |

The rationale for choosing these parameters is given as follows:

1. The number of processes in any time interval *N_p_*. This choice interacts with another free parameter, namely the number of gamma cycles used by an individual process *t_γ_*. The range of choices for both of these parameters is guided by the observation of cortical neuron spiking rates are in the 10 ~ 50 Hz range. The cycles taken by a process is reasonably guided by the distribution of gaze fixation times, typically having a mean of 200 ~ 600 milliseconds.
2. The time needed for the delay code, *d_γ_*. This choice is bounded by the observation that sensitivity to the Pulfrich pendulum illusion [Lit, 1960] that implies a sensitivity to 5 ~ 20 milliseconds and the need to keep the phase delay less than half the gamma frequency period. To this is added the refractory interval *d_r_*, taken to be 3 milliseconds.
3. The range of gamma frequencies used. This is taken to be within 30 to 80 Hz but is restricted to be in the neighborhood of 40 Hz in the simulations.
4. The number of cells (basis functions) available to represent the stimulus *n_b_*, 1000, allows multiple stimuli to be represented. Given an 50 basis functions to represent an image patch, the over-completeness factor for 4 simultaneously active computations is approximately a factor of 5.

## 3 Simulation Results

The simulation uses 1000 cells but for clarity in the visualization, a representative 50 cells are shown. The simulation time is arbitrary, but 800 milliseconds was used and 600 milliseconds is shown in the figures.

### Process mixing evaluation

The main results of the simulation are shown in Fig. 5. Owing to being generated in the context of specific gamma frequencies, individual spikes can be identified with the specific image patch that they are coding. In Fig. 5a, all of the spikes from cells that are coding the same image patch have a common color are denoting their instantaneous modulation frequency. For comparison, Fig. 10 (Supplementary Information) illustrates the simulation’s spikes as might be obtained with a conventional multi-cell recording. Without the multiprocessing context, the spike trains appear very conventional, but once the generating context is available, as shown in Fig. 5a, the spike interpretations change dramatically.

**Figure 5:**
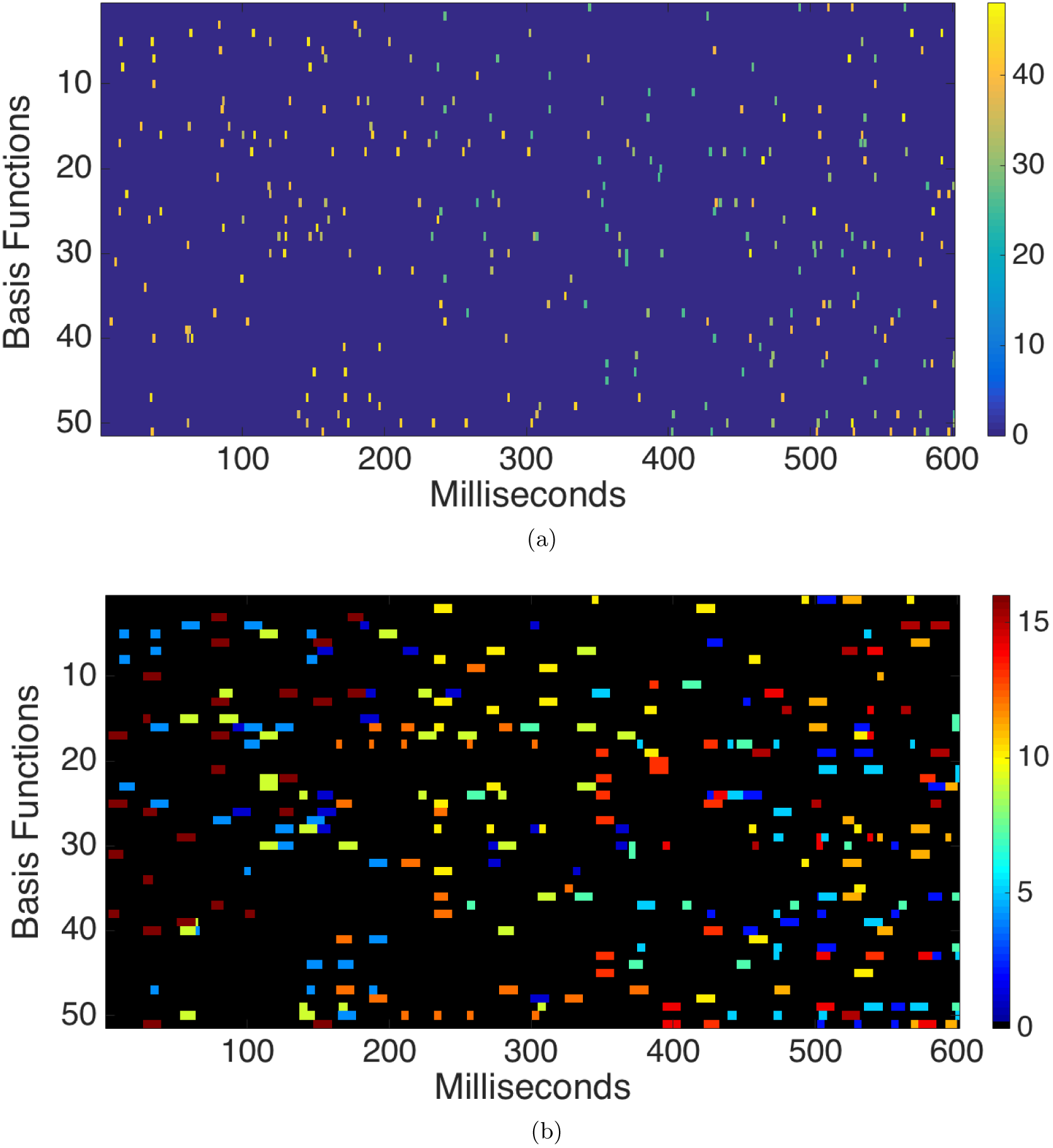
Features of the multiplexing model. The portion of the spikes produced in a larger 1000 basis function simulation with the parameters in Table 1. Spikes are rendered with 2 ms thickness for visibility. (a) The spikes in the simulations, they might appear in a simultaneous recording where rows represent specific neurons. (b) Since each spike is generated in the context of a specific gamma frequency modulator, it can be color coded by that frequency. (b)Since each spike is also part of a coding set for an individual image patch, which is here a stand-in for an arbitrary neural computational process, they can also be color coded by a particular process index in which they participate. The width of the bars indicates shows the spike generation intervals (delay+refractory) in which individual neurons are locked out from interference by other gamma modulators.

Besides the modulating frequencies there are two additional details that can be depicted. These are components of the generating processes, which utilize mechanisms that effectively prevent competing processes from using a neuron if it is in the act of generating a spike. The effects of these mechanisms is illustrated in Fig. 5b, which shows the lockout periods used in each spike’s generation. Since the simulation also keeps track of the particular process that each spike is engaged in, their particular image patch encoding, the lockout intervals can be color coded by process. Consequently each spike can be shown with an interval surround that includes its delay code plus a subsequent recovery interval.

### Modulation details

While there is a lot of room in space and time for multiplexing multiple processes, there are some subtleties discussed here. Fig. 6a makes explicit a distinctive feature of the model. The multiprocessing environment is assumed to be composed of processes that have short lifetimes and use specific gamma frequencies. The figure shows the frequencies and their temporal durations that were used to generate the corresponding spike codes. Note that the frequency band does not mean that a cell will produce a spike at any particular time while it is active, only that a cell could be instantaneously modulated at that frequency and, if selected, could produce a spike.

**Figure 6:**
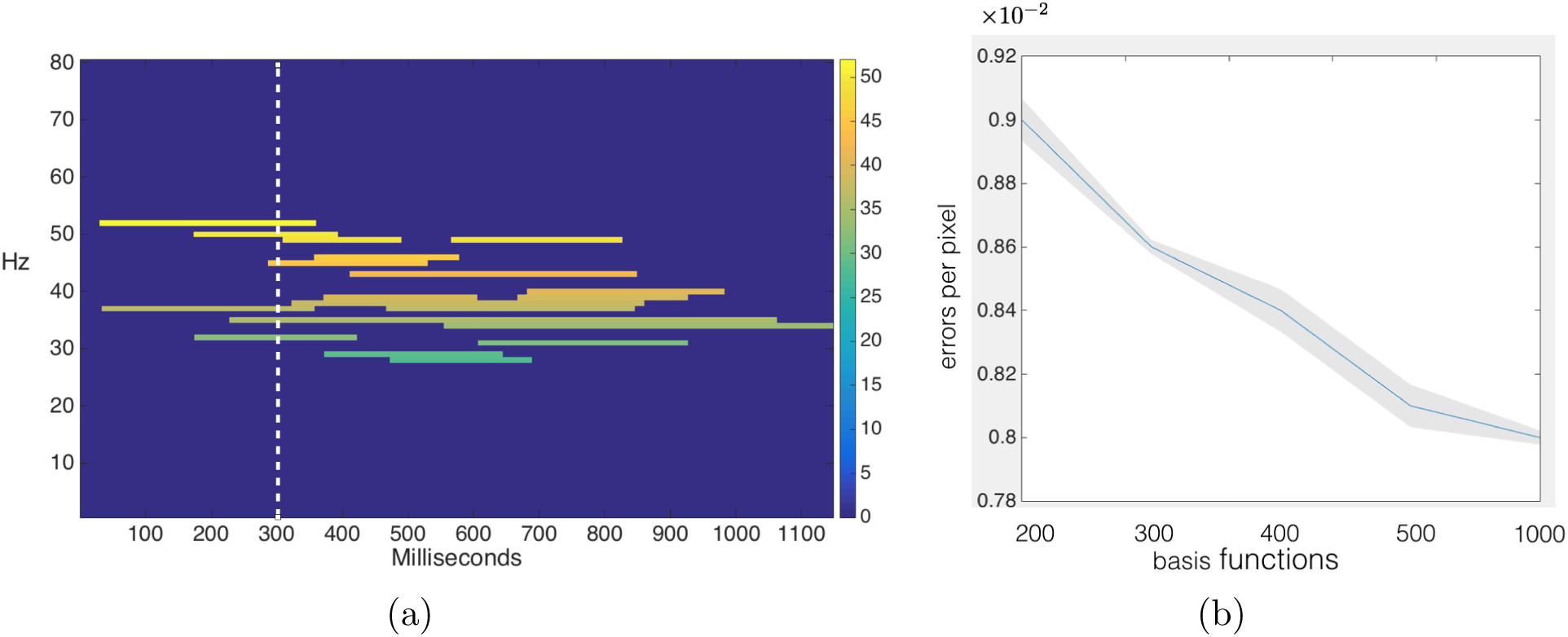
(a) The duration of a patch’s code is selected at random in the simulation and uses a specific gamma frequency selected here from the ‘low gamma’ range. The figure shows the intervals wherein the chosen frequencies are available. Consequently a prediction of the model is that on a timescale of a few hundred milliseconds the gamma spectrum should be discrete. For example at 300 milliseconds, 6 distinct gamma frequencies - 32,35,37,44,49,52 - are being co-modulated on on cell’s somas. For 52Hz, the highest frequency shown, the subset of cells that are instantaneously modulating at this frequency are available for determining the code for the associated input. (b) The coexistence of multiple gamma frequencies

The co-existence of frequencies has the consequences that all the basis functions cannot be simultaneously available for all of the frequencies. However the consequences of using less than the full complement are minor as shown by Fig. 6b, which shows that the representation accuracy degrades only slightly with less basis functions to chose from in the pool.

Another issue arises from the use of different gamma band frequencies. This situation means that when a particular frequency and input part is chosen, another frequency/input pair for a given cycle may have the same relative phase and corrupt the input. However it turns out this error source is small owing that, in the large 196-dimensional space of the input, the sparse coding strategy means that the process signals are confined to a 50-dimensional subspace. Since the axes are fairly orthogonal, the input from one input typically has a very small projection into the space of another input. If **u_a_** is a basis function for frequency *a* and **u_b_** a vector for frequency *b* that happens to have the same zero phase point, the projection onto *a*, given by 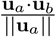 has a SEM of .007 using vectors normalized to unity. Consequently such a collision introduces a very small error for one iteration.

### Parameter sensitivity

Table 1 contains several parameters, but the most important ones are the average duration of a process in cycles, the total number of cells, the number of basis functions, and the number of processes. However in terms of generating spikes that appear Poisson distributed, the GSM model is surprisingly insensitive to these parameter settings, as shown in Fig. 7. The main reason is the random choice of the basis functions. The projections of them onto the input are very similar so they have similar probabilities of being chosen. That feature, together with the mixing effect of other processes, combine to make the spike distributions appear Poisson.

**Figure 7:**
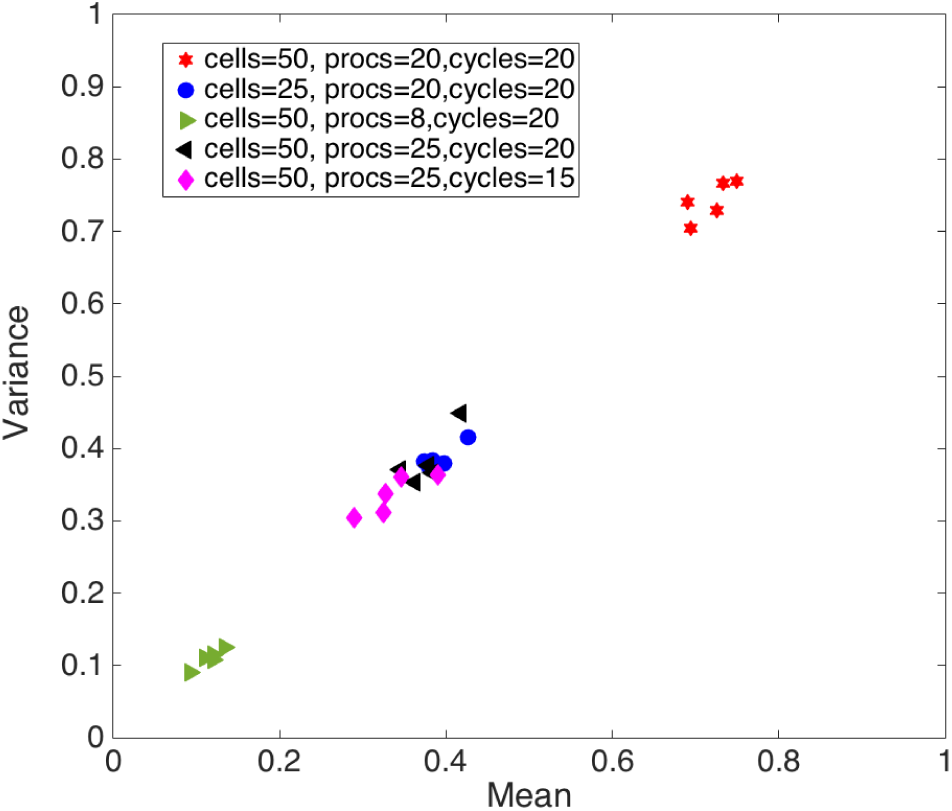
Fano ratio for different system parameters. Black squares show 5 simulations for the parameters in Table 1. In the remaining plots only the parameter that was changed from the standard set is indicated. For the red diamond, the number of cells (basis functions) was lowered to 25 instead of 50, and the the number of processes and cycles were 20 and 16 respectively. Five samples of each setting were measured. The ratios of the variance to the mean for the sames are very close to the Poisson process ideal of unity, partly because the model is noise-free.

### Trial averaging vs population averaging

Gamma frequencies are seen in some experiments and not others, and the GSM model allows the evaluation of the extent to which gamma frequencies are observable in the simulations. One distinct possibility for the differences that arise is that experimental protocols can be very different. This difference is shown in two distinct simulation data evaluation methods for evaluating the observability of the gamma signal from the model. In the classical view, data is acquired by trial averaging of a single neuron where the histories from different trials are summed once a common temporal reference is determined. The main assumption behind this is that the overall process is ergodic, that is, trial averaging is deemed to be equivalent to averaging many cells in a large network. However in the gamma multiplexing model this equivalence does not hold. The reason is that trial averaging may involve different procedures, which have different controlling parameters (such as gamma frequency or process duration) from trial to trial. In contrast, ensemble averaging uses the same control parameters for all the participating cells from a single trial). The result is that in ensemble averaging, correlations between cells over time can be evident, whereas in trial averaging, they are usually nonexistent. Fig. 8a shows the spike interval histogram for trial averaging exhibits the Poisson exponential decay with interval length. This smooth decay is much less evident in Fig. 8b, which shows the residual effects of the gamma modulation reveal themselves in periodic local maxima, as denoted by the figure’s arrows.

**Figure 8:**
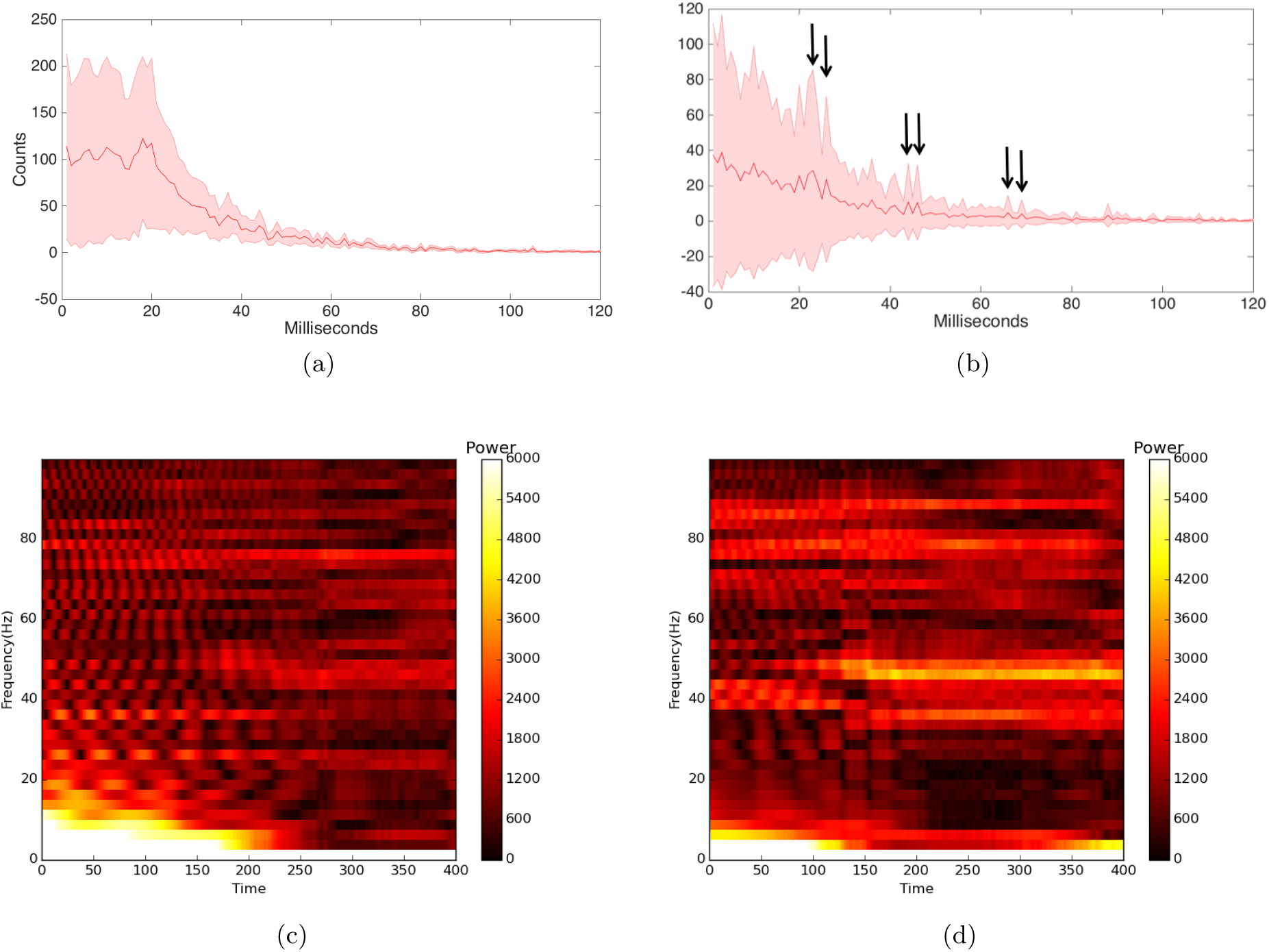
Trial averaging compared to ensemble averaging. (a) The sum of spikes interval frequencies for spikes produced in a trial averaging simulation. The characteristic decaying exponential produced by Poisson statistics is very evident. (b) The ensemble averaging histogram shows the influence of the gamma modulation (arrows). (c) Fourier spectrogram for the trial averaging cases reveals very little gamma modulation in the low gamma (35 ~ 55Hz) band owing to the fact that the controlling parameters are different for each trial. (d) The ensemble averaging Fourier spectrogram reveals marked frequency responses in the gamma band (particularly in the lower right quadrant) as each process is using a common set of parameters. Integrating the power over the {35, 55} gamma range shows that the ensemble case has 50% more power.

The computation involved in computing the interval histogram in ensemble averaging (Fig. 8b) is straightforward in that all the delays for all the basis function for the total simulation time are summed. However the method used for trial averaging takes a shortcut that approximates the desired result. Ideally one would perform 1000 simulations and then sample one of the basis function’s data to produce a histogram with a comparable number of samples. The goal is to show the effects of trial averaging using different latent parameters for each trial. To approximate this on one simulation instance the order of choices of parameters is inverted. Each basis function is chosen before the rest of parameters and thus has different sets of parameters for gamma frequency, start time and process duration. Summing intervals for the different basis functions provides a trial averaging approximation.

The difference between trial averaging and ensemble averaging is also dramatic in the spectrograms of the two cases. Figure 8c shows the spectrogram of the trial average case. In this figure we have taken the same shortcut. As the simulation shows, the gamma power is all but absent from the spectrogram. In contrast, Figure 8d shows the case where the spike averages are taken by averaging responses in an ensemble simulation. In this simulation, the gamma power is very evident.

These comparisons are driven by the multiplexing hypothesis, which obviates the assumption that the neural population in an ensemble obeys an ergodicity where repeatedly sampling one cell is equivalent to sampling a population simultaneously. The simulation shows a very evident lack of ergodicity, between these cases, but non-ergodic cases can be constructed in the ensemble averaging cases for some sets of parameters that promote mixing that appears Poisson. This property may be why gamma signals are observed in some experiments but not others.

## 4 Discussion

The model’s reconciliation of gamma and Poisson action potential observations is straightforward. The modulation of the soma at gamma rates uses the somatic potential rise time to synchronize the production of the spike. Compared to the timescale of gamma frequencies, these spikes are infrequent and thus can appear to be randomly distributed [Brunel and Wang, 2003]. This observation would be enhanced if the somatic potential modulations come in short bursts, as depicted in Fig. 1.

The focal premise of the paper is that cortical neural computation can be factored into independent processes. Although understanding the particular purpose of each individual process is very important, and though we are sympathetic to the emphasis on predictive coding in [Vinck and Bosman, 2016], an emphasis on any particular algorithm is sidestepped here to focus on the role of the neural circuitry implementing parallel communication. The functionality of a process’s subnetwork is compatible with any attractor model, which can be feed-forward, feedback, or a combination thereof. The main assumption of an attractor is that the computation needs to converge in a small number of gamma cycles.

A major issue that is sidestepped here is that of the control of a process. Multiprocessing implies that there must be a method for initiating a process and determining when it is finished. While a complete account of such a mechanism is very much a research topic, rapidly increasing evidence suggests that *α* and *β* frequencies may play this role [Michalareas et al., 2016, Van Kerkoerle et al., 2014].

Multiprocessing has not been extensively studied as an explicit focus in neuroscience, but there is ample suggestive evidence at the behavioral level that suggests the need for multiple ongoing independent neural processes. Evidence for individuated processes comes from studies of visual routines [Gilbert and Li, 2013, Roelfsema et al., 2000]. Another comes from the argument that competing motor acts are simultaneously activated [Kobak et al., 2016], implying they would co-exist as separate neural processes.

### Implications of the model for neural signaling

The gamma-latency multiplexing model provides a novel perspective on cortical computations with several new interpretations of cortical processing. Many of the features of the model have been proposed in other contexts [VanRullen and Thorpe, 2002, Womelsdorf et al., 2006, Fries et al., 2007, Landau and Fries, 2012, Bastos et al., 2012, Vinck and Bosman, 2016], but assembling them into a complete system, that incorporates, both somatic gamma modulation and probabilistic state migration results in a number of innovative features:

1. The processing speed is significantly faster than Poisson models. The speed of a basic clock cycle of a process is that of a gamma band frequency;
2. The representation is much more compact than Poisson models. Since each spike can represent an analog quantity, the circuit is not reduced to counting spikes in a unitary system;
3. The probabilistic selection is compatible with Bayesian models. Extensive evidence indicates that the brain makes extensive use of priors in interpreting data [Weiss et al., 2002, Deneve et al., 1999]. Neural processes also sample distributions in the course of computation;
4. Probabilistic selection ensures cell calibration by polling them appropriately. Probabilistic sampling ensures that all the cells in a large population have a chance of being selected and consequently having their receptive fields being updated regularly;
5. The use of delay codes is compatible with spike timing dependent plasticity (STDP) [Markram et al., 1997], although in each, implementation details need to be resolved. The delays in hippocampus [Bi and Poo, 1998] are longer than the 5 ~ 10 milliseconds used here, but that may be attributed to the use of the gamma modulating zero reference;
6. The ability to share receptive fields can account for their variability. The emerging complexity of receptive fields may be explained as the organization of receptive fields to carry multiple separate receptive subsets of synapses that are disjunctions;
7. Multi-process mixing can result in Poisson spikes in trial averaging. Superpositions of spikes are generated infrequently on different gamma cycles appear Poisson.

The proposed model is naturally sympathetic and complementary to the experimental characterizations of the gamma signal [Womelsdorf et al., 2007, Fries et al., 2007, Landau and Fries, 2012, Bastos et al., 2012, Sirota et al., 2008], but has a different interpretation of the role of the gamma frequency band. Gamma synchronization as a medium makes sense of large-scale communication; gamma synchronization in the small, addresses the organization of spike codes in the networks. The GSM model posits that its primary message transmitting role is to allow separate computations to be carried out without crosstalk, and that each coextensive process mixes the signals among neurons to foster learning and Bayesian processing. In this regard the model takes a neutral stance as to how the gamma band is sampled, but it may be that this sampling has task-specific components [Brunet et al., 2013].

### Receptive field interpretations

The traditional interpretation of neural spikes, dating from early work by Barlow Barlow [1972] is that they are engendered in service of a single process and have static interpretations. One can think of this interpretation loosely as the output being the logical ‘AND’ of its inputs. However multiplexing changes this interpretation radically in that, the output is the logical ‘OR’ of its time-coded inputs. This perspective can shed light on several experiments.

In a classic monkey experiment by Moran and Desimone, a area V4 receptive field contained two sub-fields Moran and Desimone [1985]. For discussion let us denote them A and B. By manipulating the attentional state of the animal, either one or the other sub-field was attended to, resulting in separate spike rates for each of the foci. However when the animal is attending away from the receptive field, the response is the average of the spikes for A and B. This result can be explained by multiplexing in the following way. When attending to a sub-field stimulus, the spikes for that stimulus dominate the traffic through the cell, but when attending away, the spike traffic reflects average of the separate responses to A and B, which could be produced by multiplexing. This can simply be made to happen by modulating the number of specific wavelength slots available at a cell’s soma. Attended stimuli receive more slots. This is a very different model than [Fries, 2015], wherein gamma stimuli compete in time.

This argument implies that the effects are produced distally by the representations for the sub-fields, which have their reflections in the V4 cells chosen dynamically. The notion that a cell’s observed receptive field is a consequence of distal effects is freeing in a more general way. Consider that studies of the cortical area LIP have produced evidence for its cells representation in a substantial number of very different features [Gottlieb and Snyder, 2010]. Trying to create synapses for each of these observations depends on the total synaptic budget, but there is lots of room. Taking the 14 × 14 image patch as representative and assuming a synaptic budget of 10, 000 contacts allows for 50 qualitatively different receptive fields and consequently allows a neuron to participate in that many different computations, a fraction of them simultaneously. Thus multiplexing allows many different circuits to be created for different simultaneous processes. Recent results in de-mixing spike trains are very compatible with this view [Kobak et al., 2016]. That reference shows that spikes from individual neurons can be usefully seem as mixed. In comparison to the simulations here, their data analysis shows that the majority of the spikes are seen as not interpretable as task related. In the multiplexing view additional non task-related processes are sharing the task-related cells spike generation capabilities.

### Reinterpreting attention

The model also provides an possible interpretation of the effects of attention [Maunsell, 2015, Maunsell and Treue, 2006, Boynton, 2008]. This view states that the general effects of attention can be accounted for by a gain factor. However it is not readily apparent how this advantage can be used by the perceiver. In contrast the gamma latency model suggests that the gain factor may reflect the use of additional basis functions by a computational process, that is, the size of the coding pool is increased directly resulting in additional accuracy and an increased probability of any particular neuron being included in a representation.

While attentional effects can be readily measured experimentally, nonetheless some of the effects appear as small increments on a baseline firing pattern, raising the question of how they can be reliably separated and used [Gilbert and Li, 2013, Roelfsema et al., 2000]. The GSM model has a straightforward answer to this issue in that the attentional effects are captured by separate processes that use separate frequencies and thus can be separately polled for their data. This property is particularly evident in [van Kerkoerle et al., 2016] where a large neural transient produced by a transient stimulus gap is immediately ignored afterwards on the neural recording. The ability of the neuron immediately to filter out such a large transient signal is readily understood if the transient and tracing components can be seen as separate processes that share the recorded neuron.

Another important issue is that of Bayesian models. Bayesian models have enormous support as a way the brain could use information from distal sources in reasoning e.g. [Weiss et al., 2002, Doya, 2007, Ma et al., 2006, Pouget et al., 2013b] so that there is little doubt that they are somehow implemented in neural circuitry. However an unsolved issue is the exactitude of the neural correspondence, since there is a range of possibilities between abstract models [Rao and Ballard, 1999] and some of these models that would use ensembles of spikes [Denève and Machens, 2016, Ma et al., 2006]. The gamma latency model argues for a more abstract implementation since the timed spikes can send scalar quantities.

### Caveats of the model

The model is incomplete in that important issues have been neglected in the course of the exposition of the main results. One such issue is that simulations assume that all the gamma oscillations are phase locked. While information traveling between successive neurons can be delayed, there are at least three mitigating circumstances. One is that in the substantially mylenated pyramidal cells, the spike propagation speed is sufficient to all the spike to have a margin to arrive before the next cycle. The second is that the circuitry between cortical maps may have independent couplings that allow computation to proceed in parallel. The third is that the entire cortex has only ten levels with connectivity arranged in polytrees, potentially allowing the invariant abstract maps to settle faster.

While local computation in gamma circuitry may be fast, recent evidence that shows that neurons in areas exhibit successive phase delays [Maris et al., 2016, 2013, van Ede et al., 2015]. One of the reasons for these delays would be that the cortex can speed up, in special instances, the transit of information between areas by matching phase delays with neural delays to reduce the time penalty between areas. For example phase delays could be structured to feed up the feed-forward pathway to speed up object recognition [Hung et al., 2005]. However there is no attempt to deal with this issue in the GSM simulation.

The model utilizes a simple setting of re-coding image patches from basic (LGN) representations, but a general capability required is that the new basis sets at a given zero crossing at the next gamma cycle be created from current basis sets calculated at zero crossing time. Such an algorithm has been developed [Druckmann and Chklovskii, 2010], and would be usable by the model with some adjustments to make it parallel.

In a model that depends so completely on exact timing, a very significant problem to address is that mechanisms for writing in and reading out its multiplexed messages. At the sensory end there is the problem of acquiring information from the retinal code in the first place. How can this be done? An important observation in this respect is that in its normal operation, the ubiquitous saccades and micro-saccades of primate vision naturally produce very phasic inputs. Moreover those inputs are very sensitive to delays in the range needed by the model. From this perspective, the problem reduces to one of using some kind of phase locking to place the input in register with the phase of the process. A natural site for this to happen is the thalamus, and there is evidence that such mechanisms are used in whisking [Yu et al., 2015]. At the more abstract interfaces there is the problem of dealing with the interactions with the myriad of frequencies - delta, theta, alpha and beta - that interact with the gamma band in various ways. Such interactions are the subject of extensive experimentation [Fries, 2015], but from the GSM model viewpoint, a common mechanism would seem to be some kind of phase locking between gamma and its coupled frequency. Solving this problem is likely to draw on the extensive literature studying the dynamics of such systems [Brette et al., 2007, Cannon and Kopell, 2015, Rinzel and Ermentrout, 1998].

The model assumes that once the gamma frequency modulator acquires a cell, its ensuing input can be interpreted sufficiently rapidly to produce an appropriately delayed output spike. The details of how this process could work remain to be established, but there is the possibility that the rise phase of the gamma somatic potential can sum with the dendritic input to produce appropriate spike delays.

The claims for multitasking can be seen as controversial without the appropriate context. Multitasking has been extensively studied [Pashler, 1994] and many dual-task experiments have been shown to produce a temporal overhead implying sequentiality. Similarly RSVP paradigms readily demonstrate an attentional costs for stimuli separated by less than approximately 300 milliseconds. However these situations are different from over-learned tasks, that may operate in parallel or have pre-learned transitions between sequential components, that are ubiquitous in visual processing.

A final delicate issue related to timing is that different frequencies may occasionally have the same phase zero crossing with the consequence that there is the possibility of crosstalk as separate process inputs can be summed at the same neuron. The image patch coding setting of the paper suggests an elegant solution, namely that once that the modulating frequency is determined, the likelihood that competing inputs are negligibly small. But work needs to be done to show that this property can be made to generalize.

### Testing the model

The GSM model may seem complex, but nonetheless there are several ways to test it. One most obvious way is to compare trial averaging and ensemble spectra as is done in Fig. 8. Power in the gamma range should be much more readily observed in the ensemble recordings. A very exacting test would examine the spectrum for small collections individual gamma frequencies. This would require estimating the time constants of the coextensive processes, but could be done. The expected result would be that, on a timescale of a few hundred milliseconds, the gamma spectra should be discrete, reflecting the number of independent active processes. A more direct test would be to examine whether the spikes are generated in phase with the somas’ gamma frequencies at the same time their overall distribution was Poisson. The expectation is that the relatively high gamma frequencies and low spike rates allow this relationship. Recent databases of somatic potentials [Perrenoud et al., 2016] allow this question to be addressed.

## Acknowledgement

Thanks to Ila Fiete and Alex Huk for helpful feedback on drafts of the document and thanks to Nicholas Priebe for many helpful discussions. The model benefited form being presented at the Simons program on brain and computation, particularly from Christos Papadimitriou, Fritz Sommer and Andreas Hetrz. We are also grateful to the reviewers, whose comments directed us to improve the paper substantially, with respect to both its connections to the larger modeling and experimental contexts. This work was supported by grants NIH MH 60624 and EY019174.

## Apendix: Supplementary Information

The supplementary information fleshes out secondary details that are useful for an understanding of the theoretical points. Fig. 9 shows aspects of the basis function approximations. Since the basis functions are selected probabilistically, choosing 50 from a total set of 1000, it is very unlikely that there will be many repeats. For example examining the first 16 basis functions in four separate encodings shows no repeats - even the same patch is being encoded in each instance. The second part of the figure shows repeated encodings of eight image patches. The sparse coding objective function trades off accuracy against the number of basis functions used. The 14×14 has 196 degrees of freedom, but the 50 basis functions do a reasonable job as the repeat encodings show. It can be noted that the encodings regularize the approximation, omitting some small details.

**Figure 9:**
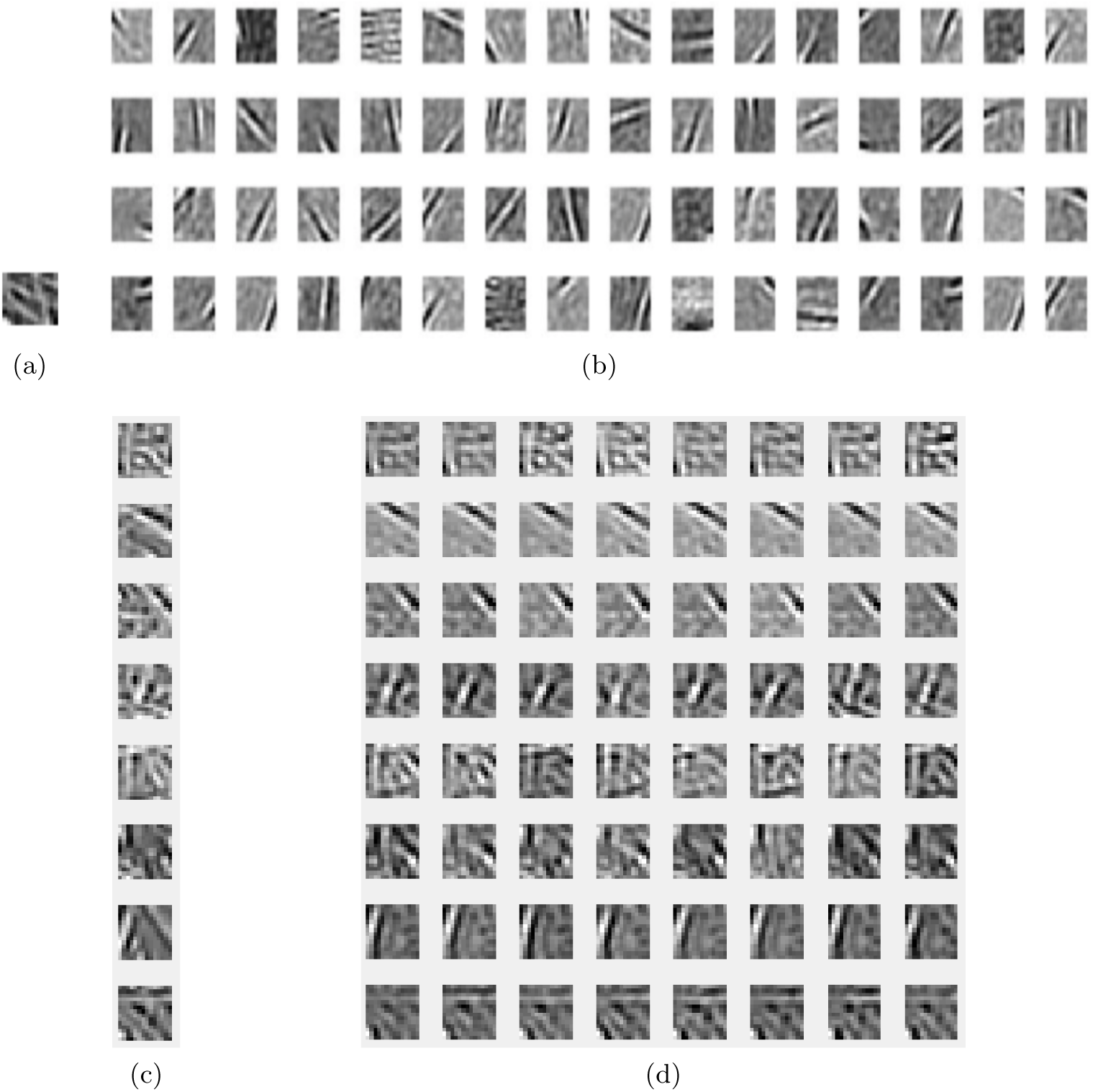
Accuracy of parallel sparse coding. (a) A single image patch of 14 × 14 pixels needs 196 degrees of freedom to be represented exactly but can be approximated by a lesser amount in an over-complete system. Here 50 cells are used. A consequence of over-completeness is that the input patch can be coded in a very large number of different ways. (b) The four rows show the first 16 cells used in four different encodings, highlighting the use of different such cells. (c,d) Eight image patches on the left reconstructed eight different times using 50 basis functions. The reconstructions are accurate with a slight tendency to regularize the patterns.

Fig. 10 shows the spikes generated in the simulation in Fig. 5 without the multiple computation indications. The assumption of the model is that this format presents information that is a correlate of a more complicate multiprocessing environment.

**Figure 10:**
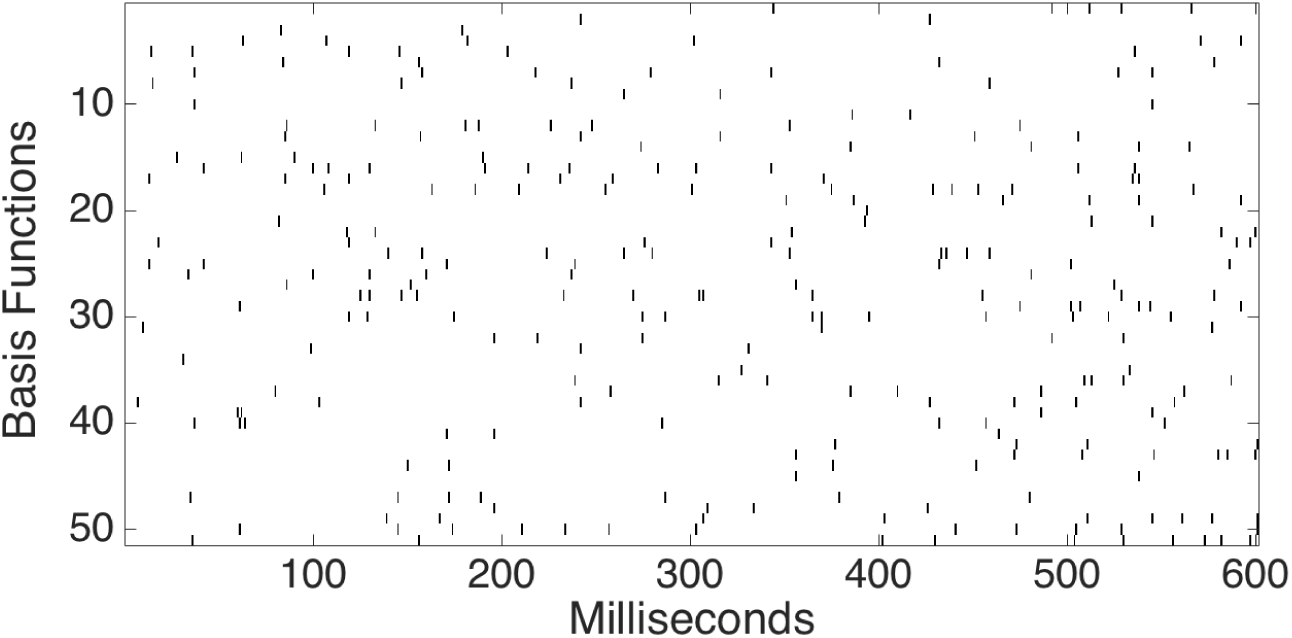
Spikes from the multiplexing model shown in Fig. 5 as they might appear in a simultaneous recording where rows represent specific neurons. Without the associated context, the spikes are extremely difficult to interpret.

A feature of the representation that is not obvious from Fig. 10 is made here in Fig. 11. For the data in the earlier figure, the spikes for each basis function are sorted by their order of generation and plotted here with the color code of the generating process. Each colored rectangle represents a single spike, showing explicitly that neurons can participate in multiple processes over time.

**Figure 11:**
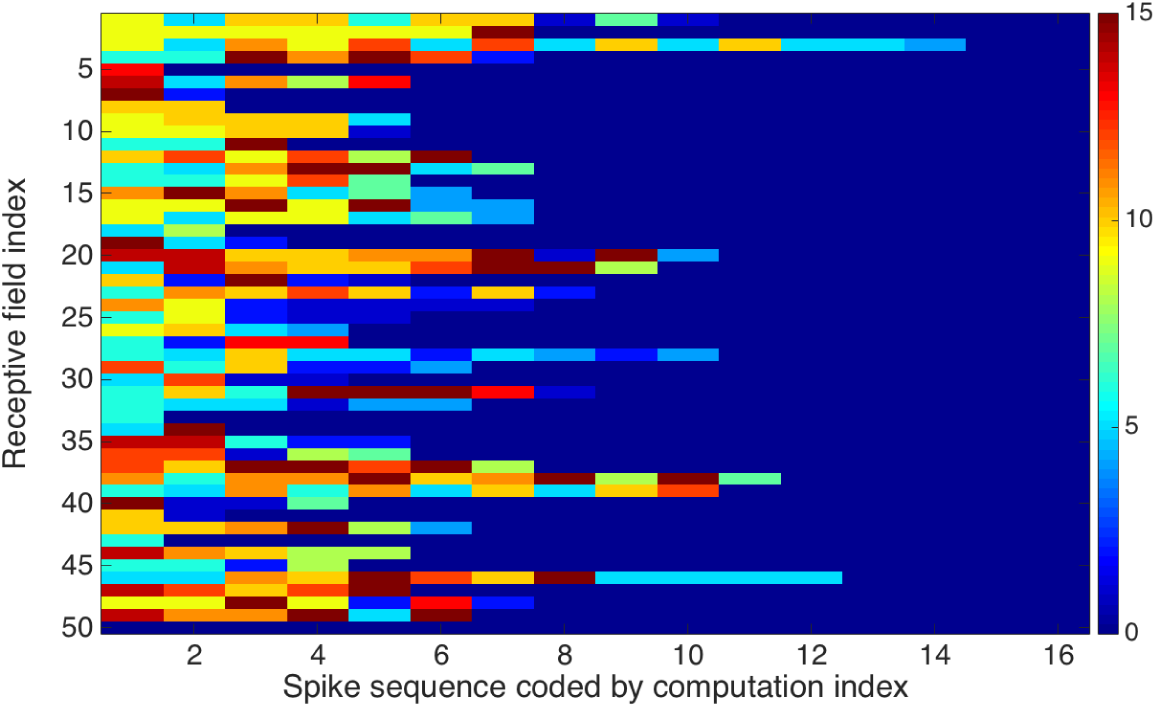
The temporal rank order of spikes in 50 neurons from the 1000 from a particular simulation. Sixteen processes are color coded allowing the multiplexing of individual neurons to be made explicit. For example reading the output for neuron 1, there are spikes from five process, the third one is uses neuron 1 again in position 5 and the forth one in position 7. Neurons tend to be reused when their receptive fields are useful in representing the input.

## Footnotes

1 Over-complete signifies that the map contains many more neurons than would mathematically be needed to represent a stimulus

## REFERENCES

Moshe Abeles. Corticonics. Cambridge University Press, 1991.

Bassam V Atallah and Massimo Scanziani. Instantaneous modulation of gamma oscillation frequency by balancing excitation with inhibition. Neuron, 62(4):566–577, 2009.

Dana Ballard and Janneke Jehee. Dual roles for spike signaling in cortical neural populations. Frontiers in computational neuroscience, 5:22, 2011.

H. B. Barlow. Single units and sensation: A neuron doctrine for perceptual psychology? Perception, 1:371–394, 1972.

Andre M. Bastos, W. Martin Usrey, Rick A. Adams and George R. Mangun, Pascal Fries, and Karl J. Friston. Canonical microcircuits for predictive coding. Neuron, 76:695–711, 2012.

J. Bernstein. Untersuchungen zur thermodynamik der bioelektrischen ströme. Pflüger’s Arch. Ges. Physiol., 92:521–562, 1902.

Michael J Berry II and Markus Meister. Refractoriness and neural precision. In Advances in Neural Information Processing Systems, pages 110–116, 1998.

Guo-qiang Bi and Mu-ming Poo. Synaptic modifications in cultured hippocampal neurons: dependence on spike timing, synaptic strength, and postsynaptic cell type. Journal of neuroscience, 18(24):10464–10472, 1998.

Geoffrey M. Boynton. A framework for describing the effects of attention on visual responses. Vision Research, 49:1129–1143, 2008.

Romain Brette, Michelle Rudolph, Ted Carnevale, Michael Hines, David Beeman, James M Bower, Markus Diesmann, Abigail Morrison, Philip H Goodman, Frederick C Harris, et al. Simulation of networks of spiking neurons: a review of tools and strategies. Journal of computational neuroscience, 23(3):349–398, 2007.

Nicolas Brunel and Xiao-Jing Wang. What determines the frequency of fast network oscillations with irregular neural discharges? i. synaptic dynamics and excitation-inhibition balance. Journal of neurophysiology, 90(1):415–430, 2003.

Nicolas Brunet, Conrado A Bosman, Mark Roberts, Robert Oostenveld, Thilo Womelsdorf, Peter De Weerd, and Pascal Fries. Visual cortical gamma-band activity during free viewing of natural images. Cerebral cortex, 25(4):918–926, 2013.

György Buzsáki and Xiao-Jing Wang. Mechanisms of gamma oscillations. Annual Review of Neuroscience, 35:203–225, 2012.

Jonathan Cannon and Nancy Kopell. The leaky oscillator: Properties of inhibition-based rhythms revealed through the singular phase response curve. SIAM Journal on Applied Dynamical Systems, 14(4):1930–1977, 2015.

Matthew Chalk, Jose L Herrero, Mark A Gieselmann, Louise S Delicato, Sascha Gotthardt, and Alexander Thiele. Attention reduces stimulus-driven gamma frequency oscillations and spike field coherence in v1. Neuron, 66:114–125, 2010.

Paul Cisek. Making decisions through a distributed consensus. Current opinion in neurobiology, 22(6):927–936, 2012.

Sophie Denève and Christian K Machens. Efficient codes and balanced networks. Nature Neuroscience, 19:375–382, 2016.

Sophie Deneve, Peter E Latham, and Alexandre Pouget. Reading population codes: a neural implementation of ideal observers. Nature neuroscience, 2(8):740–745, 1999.

A Dettner, S Münzberg, and T Tchumatchenko. Temporal pairwise spike correlations fully capture single-neuron information. Nature Communications, page 7:3805, 2016.

Markus Diesmann, Marc-Oliver Gewaltig, and Ad Aertsen. Stable propagation of synchronous spiking in cortical neural networks. Nature, 402(6761):529–533, 1999.

Kenji Doya. Bayesian brain: Probabilistic approaches to neural coding. MIT press, 2007.

Shaul Druckmann and Dmitri B Chklovskii. Over-complete representations on recurrent neural networks can support persistent percepts. In NIPS, pages 541–549, 2010.

Andreas K. Engel, Pascal Fries, Peter König, Michael Brecht, and Wolf Singer. Temporal binding, binocular rivalry, and consciousness. Consciousness and Cognition, 8:128–151, 1999.

Pascal Fries. Rhythms for cognition: communication through coherence. Neuron, 88(1):220–235, 2015.

Pascal Fries, John H Reynolds, Alan E Rorie, and Robert Desimone. Modulation of oscillatory neuronal synchronization by selective visual attention. Science, 291(5508):1560–1563, 2001.

Pascal Fries, Danko Nikolić, and Wolf Singer. The gamma cycle. Trends in neurosciences, 30(7): 309–316, 2007.

George L Gerstein and Benoit Mandelbrot. Random walk models for the spike activity of a single neuron. Biophysical journal, 4(1 Pt 1):41, 1964.

Wulfram Gerstner, Andreas K Kreiter, Henry Markram, and Andreas VM Herz. Neural codes: firing rates and beyond. Proceedings of the National Academy of Sciences, 94(24):12740–12741, 1997.

Charles D Gilbert and Wu Li. Top-down influences on visual processing. Nature Reviews Neuroscience, 14(5):350–363, 2013.

Jacqueline Gottlieb and Lawrence H Snyder. Spatial and non-spatial functions of the parietal cortex. Current opinion in neurobiology, 20:731–740, 2010.

Charles M Gray, Peter König, Andreas K Engel, Wolf Singer, et al. Oscillatory responses in cat visual cortex exhibit inter-columnar synchronization which reflects global stimulus properties. Nature, 338(6213):334–337, 1989.

Ann M. Graybiel, Toshihiko Aosaki, Alice W. Flaherty, and Minoru Kimura. The basal ganglia and adaptive motor control. Science, 365:1826–1831, 1994.

Stephen M Highstein, Richard D Rabbitt, Gay R Holstein, and Richard D Boyle. Determinants of spatial and temporal coding by semicircular canal afferents. Journal of neurophysiology, 93(5): 2359–2370, 2005.

A. L. Hodgkin and A. F. Huxley. A quantitative description of membrane current and its application to conduction and excitation in nerve. Journal of Physiology, 117:500–544, 1952.

Chou P Hung, Gabriel Kreiman, Tomaso Poggio, and James J DiCarlo. Fast readout of object identity from macaque inferior temporal cortex. Science, 310(5749):863–866, 2005.

Janneke FM Jehee and Dana H Ballard. Predictive feedback can account for biphasic responses in the lateral geniculate nucleus. PLoS Comput Biol, 5(5):e1000373, 2009.

Janneke FM Jehee, Constantin Rothkopf, Jeffrey M Beck, and Dana H Ballard. Learning receptive fields using predictive feedback. Journal of Physiology-Paris, 100(1):125–132, 2006a.

Janneke FM Jehee, Constantin Rothkopf, Jeffrey M Beck, and Dana H Ballard. Learning receptive fields using predictive feedback. Journal of Physiology-Paris, 100(1):125–132, 2006b.

Dmitry Kobak, Wieland Brendel, Christos Constantinidis, Claudia E Feierstein, Adam Kepecs, Zachary F Mainen, Ranulfo Romo, Xue-Lian Qi, Naoshige Uchida, and Christian K Machens. Demixed principal component analysis of neural population data. Elife, 5:e10989, 2016.

Ayelet Nina Landau and Pascal Fries. Attention samples stimuli rhythmically. Current Biology, 22:1000–1004, 2012.

P. Lennie. The cost of cortical computation. Current Biology, 13:493, 2003.

Alfred Lit. The magnitude of the pulfrich stereo-phenomenum as a function of target velocity. Journal of Experimental psychology, 59:165–175, 1960.

W. J. Ma, J. M. Beck, P. E. Latham, and A. Pouget. Bayesian inference with probabilistic population codes. Nature Neurocience, 9:1432–1438, 2006.

Eric Maris, Thilo Womelsdorf, Robert Desimone, and Pascal Fries. Rhythmic neuronal synchronization in visual cortex entails spatial phase relation diversity that is modulated by stimulation and attention. Neuroimage, 74:99–116, 2013.

Eric Maris, Pascal Fries, and Freek van Ede. Diverse phase relations among neuronal rhythms and their potential function. Trends in neurosciences, 39(2):86–99, 2016.

Henry Markram, Joachim Lübke, Michael Frotscher, and Bert Sakmann. Regulation of synaptic efficacy by coincidence of postsynaptic aps and epsps. Science, 275(5297):213–215, 1997.

JohnH.R. Maunsell. Neuronal mechanisms of visual attention. Annual Review of Vison Science, 1:373–391, 2015.

John H.R. Maunsell and Stefan Treue. Feature-based attention in visual cortex. TRENDS in Neurosciences, 29:317–322, 2006.

Georgios Michalareas, Julien Vezoli, Stan Van Pelt, Jan-Mathijs Schoffelen, Henry Kennedy, and Pascal Fries. Alpha-beta and gamma rhythms subserve feedback and feedforward influences among human visual cortical areas. Neuron, 89(2):384–397, 2016.

Jeffrey Moran and Robert Desimone. Selective attention gates visual processing in the extrastriate cortex. Frontiers in cognitive neuroscience, 229:342–345, 1985.

B. A. Olshausen and D. J. Field. Sparse coding with an overcomplete basis set: A strategy employed by v1? Vision Research, 37:3311–3325, 1997.

Harold Pashler. Dual-task interference in simple tasks: data and theory. Psychological bulletin, 116:220, 1994.

Judea Pearl. Fusion, propagation, and structuring in belief networks. Artificial intelligence, 29(3): 241–288, 1986.

Judea Pearl. Probabilistic reasoning in intelligent systems: networks of plausible inference. Morgan Kaufmann, 2014.

Quentin Perrenoud, Cyriel MA Pennartz, and Luc J Gentet. Membrane potential dynamics of spontaneous and visually evoked gamma activity in v1 of awake mice. PLoS biology, 14(2): e1002383, 2016.

Jean-Pascal Pfister and Wulfram Gerstner. Triplets of spikes in a model of spike timing-dependent plasticity. Journal of Neuroscience, 26(38):9673–9682, 2006.

Jonathan W Pillow, Liam Paninski, Valerie J Uzzell, Eero P Simoncelli, and EJ Chichilnisky. Prediction and decoding of retinal ganglion cell responses with a probabilistic spiking model. Journal of Neuroscience, 25(47):11003–11013, 2005.

Alexandre Pouget, Jeffrey M Beck, Wei Ji Ma, and Peter E Latham. Probabilistic brains: knowns and unknowns. Nature neuroscience, 16(9):1170–1178, 2013a.

Alexandre Pouget, Jeffrey M Beck, Wei Ji Ma, and Peter E Latham. Probabilistic brains: knowns and unknowns. Nature Reviews Neurocience, 16:1170–1178, 2013b.

Rajesh P. N. Rao and Dana H. Ballard. Predictive coding in the visual cortex: a functional interpretation of some extra-classical receptive field effects. Nature Neuroscience, 2:79–87, 1999.

John Rinzel and G Bard Ermentrout. Analysis of neural excitability and oscillations. Methods in neuronal modeling, 2:251–292, 1998.

Pieter R. Roelfsema, Victor A.F. Lamme, and Henk Spekreijse. The implementation of visual routines. Vision research, 40:1385–1411, 2000.

Almut Schüz and Günther Palm. Density of neurons and synapses in the cerebral cortex of the mouse. Journal of Comparative Neurology, 286(4):442–455, 1989.

Kiley Seymour, Colin WG Clifford, Nikos K Logothetis, and Andreas Bartels. The coding of color, motion, and their conjunction in the human visual cortex. Current Biology, 19(3):177–183, 2009.

M. N. Shadlen and W. T. Newsome. The variable discharge of cortical neurons: Implications for connectivity, computation, and information processing. Journal of Neuroscience, 18:3870–3896, 1998.

W Singer. Neuronal synchrony: A versatile code review for the definition of relations? Neuron, 24 (24):49–64, 1999a.

Wolf Singer. Neuronal synchrony: A versatile code review for the definition of relations? Neuron, 24:49–65, 1999b.

Anton Sirota, Sean Montgomery, Shigeyoshi Fujisawa, Yoshikazu Isomura, Michael Zugaro, and Gÿorgy Buzśaki. Entrainment of neocortical neurons and gamma oscillations by the hippocampal theta rhythm. Neuron, 60(4):683–697, 2008.

William E Skaggs and Bruce L McNaughton. Theta phase precession in hippocampal. Hippocampus, 6:149–172, 1996.

Sameet Sreenivasan and Ila Fiete. Grid cells generate an analog error-correcting code for singularly precise neural computation. Nature neuroscience, 14(10):1330–1337, 2011.

Freek van Ede, Stan Van Pelt, Pascal Fries, and Eric Maris. Both ongoing alpha and visually induced gamma oscillations show reliable diversity in their across-site phase-relations. Journal of neurophysiology, 113(5):1556–1563, 2015.

Timo Van Kerkoerle, Matthew W Self, Bruno Dagnino, Marie-Alice Gariel-Mathis, Jasper Poort, Chris Van Der Togt, and Pieter R Roelfsema. Alpha and gamma oscillations characterize feedback and feedforward processing in monkey visual cortex. Proceedings of the National Academy of Sciences, 111(40):14332–14341, 2014.

Timo van Kerkoerle, Matthew W. Self, and Pieter R. Roelfsema. The influence of attention and working memory on neuronal activity in the different layers of primary visual cortex. Nature Communications, 2016.

R. VanRullen and S. J. Thorpe. Surfing a spike wave down the ventral stream. Vision Research, 42:2593, 2002.

Martin Vinck and Conrado A Bosman. More gamma more predictions: gamma-synchronization as a key mechanism for efficient integration of classical receptive field inputs with surround predictions. Frontiers in systems neuroscience, 10, 2016.

Martin Vinck, Bruss Lima, Thilo Womelsdorf, Robert Oostenveld, Wolf Singer, Sergio Neuen-schwander, and Pascal Fries. Gamma-phase shifting in awake monkey visual cortex. Journal of Neuroscience, 30(4):1250–1257, 2010.

Christoph Von der Malsburg. The what and why of binding: the modelers perspective. Neuron, 24 (1):95–104, 1999.

Astrid Von Stein and Johannes Sarnthein. Different frequencies for different scales of cortical integration: from local gamma to long range alpha/theta synchronization. International journal of psychophysiology, 38(3):301–313, 2000.

Yair Weiss, Eero P. Simoncelli, and Edward H. Adelson. Motion illusions as optimal percepts. Nature Neurocience, 5:598–604, 2002.

Thilo Womelsdorf, Pascal Fries, Partha P Mitra, and Robert Desimone. Gamma-band synchronization in visual cortex predicts speed of change detection. Nature, 439(7077):733–736, 2006.

Thilo Womelsdorf, Jan-Mathijs Schoffelen, Robert Oostenveld, Wolf Singer, Robert Desimone, Andreas K. Engel, and Pascal Fries. Modulation of neuronal interactions through neuronal synchronization. Science, 316:1610–1612, 2007.

C. Yu, G. Horev, N. Rubinand D. Derdikman, and and E. Ahissar S. Haidarliu. Coding of object location in the vibrissal thalamocortical system. Cerebral Cortex, 25(563–577), 2015.

